# Site-specific validation and quantification of RNA 2′-O-methylation by qPCR with RNase H

**DOI:** 10.1101/2022.05.23.493005

**Authors:** Yifan Wu, Yao Tang, Yong Li, Xiangwen Gu, Qiang Wang, Qihan Chen

## Abstract

RNA 2′-O-methylation, one of the most abundant modifications on RNAs, is crucial for diverse intracellular biological processes. In the past several years, several high-throughput screening methods have been developed, resulting in the identification of thousands of new 2′-O-methylation (Nm) sites. However, due to the high variability in these high-throughput methods, accurate and rapid low-throughput validation assays are needed to confirm and quantify the 2′-O-methylation status of screened candidate sites. Although several low-throughput Nm site detection methods have been reported, precise location and quantitative assays are still challenging to achieve. Based on the characteristic that RNase H would be inhibited by Nm modification, we developed Nm-VAQ (site-specific 2′-O-methylation (Nm) <u>V</u>alidation and <u>A</u>bsolute <u>Q</u>uantification resolution). In this study, with multiple tests of reagents and conditions, Nm-VAQ was established with a chimera probe of RNA/DNA, RNase H site-specific cleavage, and qRT-PCR, which demonstrated precise absolute quantification of modification ratios and methylation copy numbers. With the help of Nm-VAQ, the 2′-O-methylation status of 5 sites in rRNA was evaluated.

## INTRODUCTION

RNA chemical modifications are pivotal for post-transcriptional regulation of gene expression. Among these, 2′-O-methylation is one most abundant modification occurring on the 2′-hydroxyl group of ribose and is present in all major classes of RNA, including rRNA, tRNA, miRNA, and mRNA (1,2). Since 2′-O-methylation is not a base-limited modification, it is called Nm (N refers to A/G/C/U)(3). Based on previous studies, Nm is commonly distributed within conserved regions of rRNA and influences rRNA folding, assembly, and metabolism by enhancing hydrophobic surfaces and stabilizing helical stem structures(3). In addition, Nm regulates various biological processes by affecting RNA-RNA and RNA-protein interactions, including splicing, degradation, translation, and immune recognition(4-7). Thus, 2-O-methyltransferases and changes to Nm levels are linked to many diseases, including cancers, autoimmune diseases, and intellectual disability (Genes (Basel) 2019 10(20):117). Given the significance of Nm modifications, the complete distribution map and regulation mechanism of Nm in different biological contexts warrant further elucidation.

To further characterize RNA 2′-O-methylation function, several high-throughput Nm identification tools have been established, such as 2′OMe-seq, RiboMethSeq, Nm-Seq, and NJU-seq(8-11). However, although these tools can detect potential Nm sites comprehensively, the results’ ambiguity leads to the significant difficulties of Nm quantification due to potential false positive and semi-quantification results among the various methods (12-14). Therefore, an accurate method is required to validate and quantify the Nm sites detected through the high throughput methods. We developed an RNase H-based site-specific 2′-O-methylation (Nm) <u>V</u>alidation and <u>A</u>bsolute <u>Q</u>uantification resolution (Nm-VAQ) protocol to address this problem. RNase H is a non-sequence-specific endonuclease enzyme that catalyzes the cleavage of RNA in RNA/DNA substrates, but its activity is inhibited by 2′-O-methylated residues(15). In previous studies, researchers had tried to achieve site-specific cleavage of RNase H by the guidance of an RNA-DNA chimera probe to evaluate potential Nm sites(15). However, the conclusion of which probe was capable seemed not consistent (15-17). Here, by testing multiple designs and continuously improving and optimizing, we established Nm-VAQ by combining RNase H cleavage property and qRT-PCR assay to acquire the absolute quantification of methylation copy number and the 2′-O-methylation ratio of the target site (Figure 1). We used Nm-VAQ to evaluate five sites in rRNA of the HeLa cell line, including 18s 159A, 354U, and 1391C (known Nm sites reported in the previous studies), 28s 4109C (newly discovered Nm site in our recent research), and 18s 1197G (unmethylated sites reported in previous studies) (11,18).

**Figure 1.**
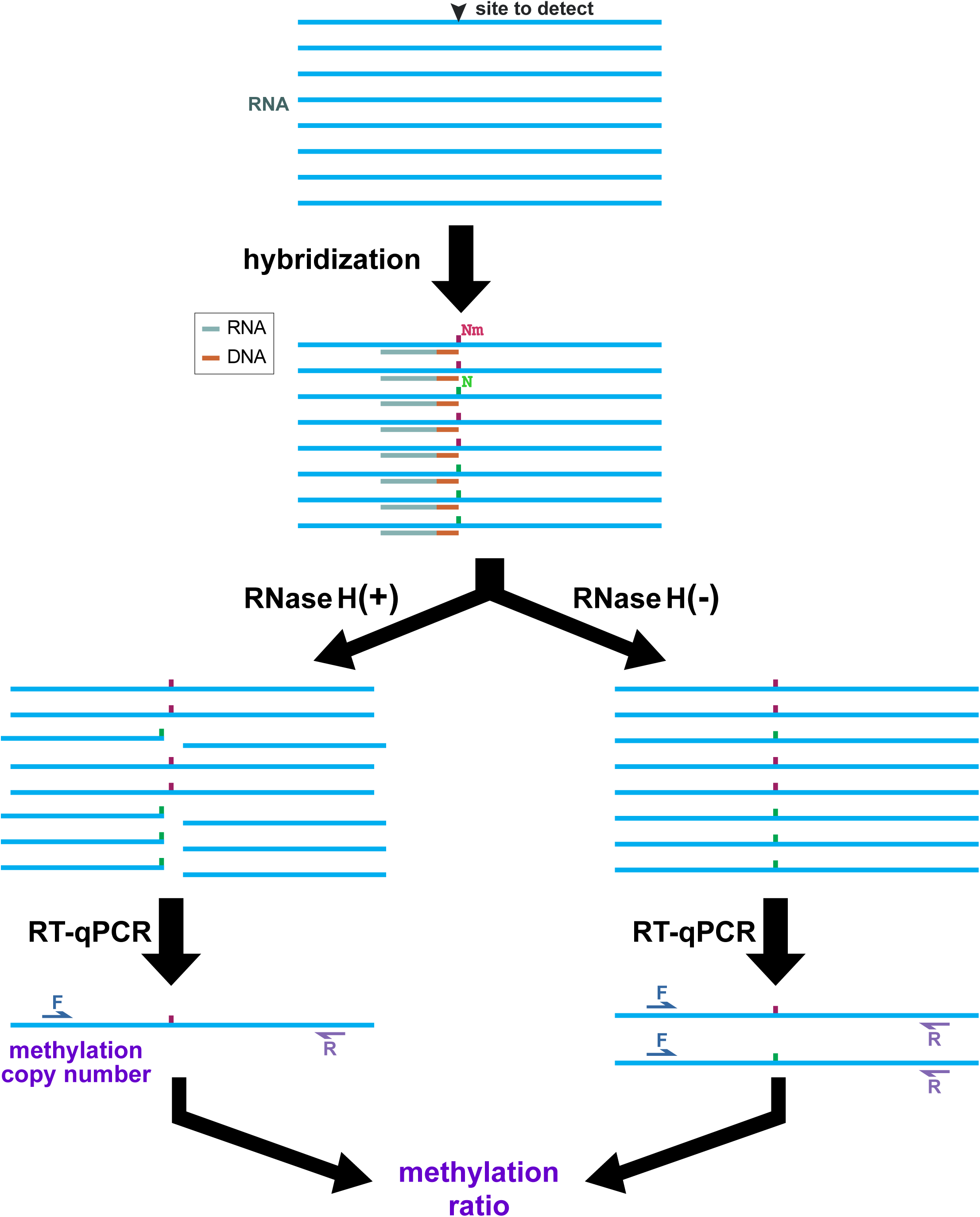
Schematic workflow of Nm-VAQ. 1) the hybrid of RNA and chimera probes. The red site in substrate indicated the target site was 2′ -O-methylated, and the green site indicated the unmethylated site. The reddish-brown region of chimera probe showed the DNA, and the baby blue region showed the RNA with Nm modification; 2) with/without RNase H cleavage; 3) RT-qPCR. Methylation copy number was calculated by CT (Cycle Threshold) of RNase H reaction. Methylation ratio was from ΔCT of RNase H cleavage and control sample. The box on the right showed an example of RNase H cleavage directed by the chimera probe in the following tests.

## MATERIALS AND METHODS

### The assay of RNase H cleavage

GenScript Biotech Co. synthesized RNA oligonucleotides and chimera probes. The sequences of all probes used in this study are listed in Figure 2 and Supplementary Figure 1. Briefly, 12.5 pmol RNA oligonucleotides were mixed with 75 pmol chimera probe, and then heated to 95 °C for 2 min, then cooling to 22 °C at 0.1 °C/s, and maintained for 5 min. The hybrid was reacted with 1 μl RNase H (New England BioLabs) at 37 °C for 30 min and then heated to 90 °C for 10 min to terminate the reaction. The cleavage products were added to RNA Dye (New England BioLabs) and then were analyzed by 20% UERA-PAGE, visualizing by ChemiDoc XRS+\UnUniversal alHoodII gel imaging system (Bio-rad). Series reactions were designed to test the duration time, cooling rate, and molar number ratio of oligonucleotides and chimera probe to improve RNase H-dependent Nm detection.

**Figure 2.**
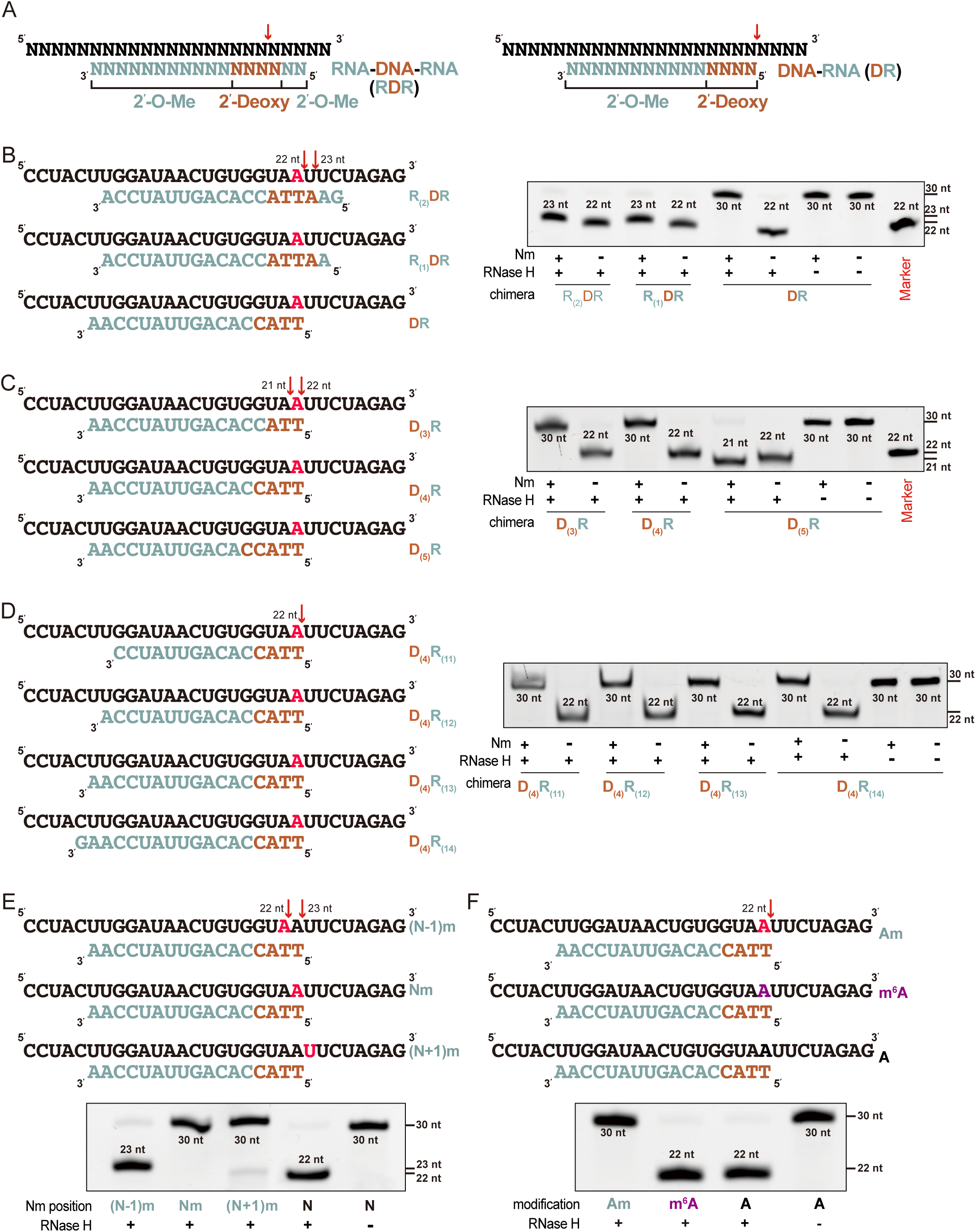
Screening chimera probe for RNase H-dependent Nm detection. (A) Schematic presentation of hybrid of RNA oligonucleotides (up sequences) and chimera probes (down sequences). The red arrow indicated the RNase H cleavage on the previous report(15,17). The baby blue sites indicated RNA of chimera probes, which all were 2′-O-methylated, and the reddish-brown sites indicated the DNA of chimera. The following chimeras were labeled in the same way. (B-F) The exploration of chimera probe structure. The scheme of hybrid RNA oligonucleotides (up sequences) and chimera probes (down sequences) was shown on the left. The RNase H reaction products were presented on the right by electrophoresis. The red sites in the substrates indicated the Nm site, and the red arrows indicated the cleavage sites. The number presented the length of FAM-labeled cleavage products. (B) Site-specific cleavage of RNase H directed by RDR or DR chimera. (C) RNase H cleavage directed by DR chimera with varied DNAs. (D) RNase H cleavage directed by DR chimera with varied RNA numbers. (E) Effect of 2′-O-methylation positions on RNase H cleavage. N represented the target site. (F) Effect of m^6^A and Nm modification on RNase H cleavage. The purple site was an m^6^A-modified site.

### Cell culture and RNA extraction

The HeLa cells were obtained from Shanghai Institute of Cell Biology, Chinese Academy of Sciences (Shanghai, China). The HeLa cells were grown in Dulbecco’ s modified eagle medium (DMEMs) supplemented with 10% fetal bovine serum (FBS), 5% antibiotic at 37 °C in an incubator containing 5% CO_2_. The cells were lysed by TRIzol (Invitrogen), and total RNA was extracted following standard protocol. The RNA amount was quantified by NanoDrop (Thermo Fisher Scientific).

### Nm-VAQ assay of RNA 2′-O-methylation ratio and copy numbers

GenScript Biotech Co. synthesized synthetic RNA oligonucleotides with/without Nm, and Sangon Biotech Co. synthesized qRT-PCR primers. All sequences are listed in Supplementary Table 1. The oligonucleotides with/without Nm were mixed to obtain gradient 2′-O-methylation ratio substrate. Then the substrate was added to 10 pmol D_(4)_R_(13)_ chimera probe, with 95 °C for 2 min, following by cooling to 22 °C at 0.1 °C/s, and 22 °C for 5 min. The products were divided into 2 parts for the following reaction. One mixture contained 5 μl previous product, 1 μl RNase H (New England BioLabs), 1 μl 10X RNase H Reaction Buffer, and 3 μl RNase-free H_2_O. RNase H storage buffer was substituted for RNase H to form the other mixture to serve as a blank control, followed by 30 min at 37 °C and 10 min at 90 °C. The products were diluted for 50-folds and then were used to obtain cDNA by HiScript II 1st Strand cDNA Synthesis Kit (Vazyme biotech co., ltd.). cDNA was diluted for 100-folds, and then qPCR was conducted in 20 μl reaction mixture containing 10 μl ChamQ Universal SYBR qPCR Master Mix (Vazyme biotech co., ltd.), 0.4 μl F primer, 0.4 μl R primer, 2 μl cDNA, and 7.2 μl H_2_O, following the protocol: 30s at 95 °C, then 40 cycles of 95 °C for 10s and 60°C for 30s. Each cDNA was analyzed in 3 replicates. Nm ratio was calculated by ΔCT (Cycle Threshold) of RNase H reaction and control, and CT of RNase H reaction obtained methylation copy number.

### Quantitation of HeLa rRNA Nm ratio by Nm-VAQ

All primers were obtained from Sangon Biotech Co, and the sequences were shown in Supplementary Table 2. 100 ng HeLa RNA and 10 pmol chimera probes were hybridized, followed by RNase H cleavage, reverse transcription, and qPCR according to the above Nm-VAQ protocol. The qPCR protocol changed the Nm detection of 18S rRNA 1391C sites with an extension temperature of 55°C in the qPCR protocol. ΔCT calculated the ratio of modification according to the Nm-VAQ standard curve.

### Quantitation of HeLa rRNA Nm ratio by RTL-P method

Two sets of primers were designed for the site, with the Fu forward primer located upstream of the Nm site and the Fd forward primer located downstream of the Nm site. All the sequences were shown in Supplementary Table 3. The high dNTPs concentration reaction mixture consisted of 5× RT Buffer 4 μl, M-MLV (H-) Reverse Transcriptase (200 U/μL) 1μl, RNase inhibitor (40 U/μL) 1μl, RNA 100 ng, RT primer (10 μM) 1 μl, 8 μl RNase -free H_2_O, dNTPs (1 mM each) 1 μl. The low dNTPs concentration reaction mixture was replaced by dNTPs (2 μM each). Then the mixtures were incubated at 45 °C for 1 hour and 85 °C for 2 min. The cDNA was diluted 100-fold and subsequently subjected to qPCR reactions, with both Fu and Fd products amplified for each cDNA. RT efficiency and RT fold change were calculated according to the following strategy. RT efficiency=template amount measured by Fu and R/template amount measured by Fd and R. RT fold change=RT efficiency with low dNTPs/RT efficiency with high dNTPs(14).

## Results

### Screening guide chimera with anchored cleavage sites at the Nm residues

Previously, Inoue and Lapham proposed two types of RNA-DNA chimera to apply site-specific cleavage of target RNA, RNA-DNA-RNA (RDR) and DNA-RNA (DR)(16,17), and all RNA of chimera were 2′-O-methylated to improve stability (Figure 2A).

We started from different chimera structures to determine which one anchors the cleavage at the targeted Nm site. A pair of synthesized FAM-labeled 30nt ssRNAs, which contained 2′-O-methylated/unmethylated A at position 22nt, were used as substrates (Figure 2B). First, we adjusted the number of RNA of the 5′ end of chimera probes, including R_(2)_DR (with two ribonucleotides at 5′ end), R_(1)_DR (with one ribonucleotide at 5′ end). Substrates were gently hybridized with the chimera at slowly cooling temperature to form a hybrid strand, then incubated with RNase H and detected by electrophoresis (see Methods). Although both R_(1)_DR and R_(2)_DR chimera probes induced specific cleavage sites on the unmethylated substrate and produced 22nt products, they were not inhibited by 2′-O-methylation completely, resulting in the production of 23nt cleavage products as well (Figure 2B), which seemed not consistent with previous studies (15-17). On the other hand, a design of chimera probe with only DR (without ribonucleotide at 5′ end) demonstrated clear site-specific cleavage on the unmethylated substrate but not 2′-O-methylation substrates. Similar results were also shown in the cleavage of another two RNA substrates (Supplementary Figure 1). Therefore, the DR structure was chosen for further modifications in subsequent experiments. The next question was whether the length of the deoxyribonucleotide part determines the cleavage site since this conclusion was not consistent in previous studies. Based on the results of D_(3)_R to D_(5)_R, cleavage activity of probes D_(3)_R and D_(4)_R were completely inhibited by 2′-O-methylation (Figure 2C). Due to the higher stability of DNA over RNA, D_(4)_R was chosen for subsequent testing. In addition, the length of the ribonucleotide part was also essential to determine the binding specificity and affinity of the DR chimera probe. To optimize the best length, we performed a similar test from D_(4)_R_(11)_ to D_(4)_R_(14)_. As shown in Figure 2D, there was no significant difference in cleavage sites or cleavage efficiency among tests with different chimera probes. In principle, the longer length of R required a higher melting temperature, facilitating the test of Nm sites in strong secondary structural regions.

### RNase H-dependent Nm detection with site-specificity and Nm-modification specificity

To further evaluate the effect of Nm around the cleavage site, we applied the assay with RNA substrate with Nm positioned on 1 nt downstream or upstream of the target ribonucleotide. As shown in Figure 2E, RNase H activity was inhibited by 2′-O-methylation of the target and 1 nt downstream ribonucleotide but not the upstream one. Thus, the combination use of chimera probes of targeting adjacent ribonucleotides can locate the accurate position of the Nm site.

So far, more than a hundred RNA modifications have been identified, most of which occurred on bases. To further confirm RNase H cleavage activity was only sensitive to Nm, we tested m^6^A, another widely distributed RNA modification, with DR chimera probe and RNase H. As expected, the m^6^A-containing substrate was cleaved by RNase H at the modification site, while Am inhibited the cleavage completely (Figure 2F).

### Improved RNase H-dependent Nm detection

We optimized the conditions since RNase H cleavage is crucial for discriminating 2′-O-methylated RNA from unmethylated RNA molecules. To achieve the full substrate-probe hybrid at the first and also critical step, 10 times more probe was added to the reaction, followed by a denaturing step of 95 °C for 2 min and a slow cooling step from 95 °C to 22 °C at 0.1 °C /s. The substrates were cleaved entirely in the different denaturation time gradients, indicating that these treatments were sufficient (Supplementary Figure 2A). To avoid RNA being damaged under prolonged high temperatures, we chose denaturation at 95 °C for 2 min to form a hybridization duplex. After treatments with the same denaturation condition, all slow cooling from 95 °C to 22 °C allows RNase H to hydrolyze completely (Supplementary Figure 2B). In addition, the molar ratio of RNA substrate and chimera probe was also tested. When the molar ratio reached 1:1, RNase H’ s substrates were wholly digested (Supplementary Figure 2C). As the more complex structure of RNA in biological samples, the slow cooling of 0.1 °C /s and the 1:10 ratio of substrate and probe were chosen for subsequent analysis.

### Construction of Nm-VAQ, a tool for Nm quantitative detection

Although several Nm detection and validation methods have been reported previously, the Nm quantitative detection is still challenging to achieve, especially on low-content RNAs or low-modified sites. To establish an accurate quantification tool, we applied RNase H cleavage directed by chimera and qRT-PCR combination, named Nm-VAQ (Nm <u>V</u>alidation and <u>A</u>bsolute <u>Q</u>uantification method).

Synthetic RNA oligos used in the previous tests were mixed with multiple ratios to assess whether Nm-VAQ can effectively determine the Nm ratio on partially methylated sites. As shown in the schematic (Figure 1), the sample was divided into two parts to incubate with/without RNase H after forming the RNA-chimera hybrid. The total target RNA copy number can be calculated by Ct value (Cycle threshold) without RNase H treatment, while the 2′-O-methylation ratio can be acquired from the ΔCT of two reactions. A highly correlated linear curve of 2′-O-methylation ratio and ΔCT was obtained (R^2^>0.99, Linear Regression Analysis), indicating that Nm-VAQ can quantify the Nm ratio accurately (Figure 3A). Although most previous Nm quantification methods demonstrated good performance on the Synthetic RNA, one of the most challenging points was the unknown amount/concentration of target RNA, which varied the result. We tested 50% Nm ratio substrate with 1 pmol, 0.1 pmol, 0.01 pmol, and 0.001 pmol concentration. Nm-VAQ demonstrated consistent results with no significant difference around 50% (Figure 3B). Furthermore, the copy numbers of substrates seemed quite linear after RNase H cleavage, which proved that RNase H would not cleave 2′-O-methylated substrates even at shallow concentrations (Figure 3C).

**Figure 3.**
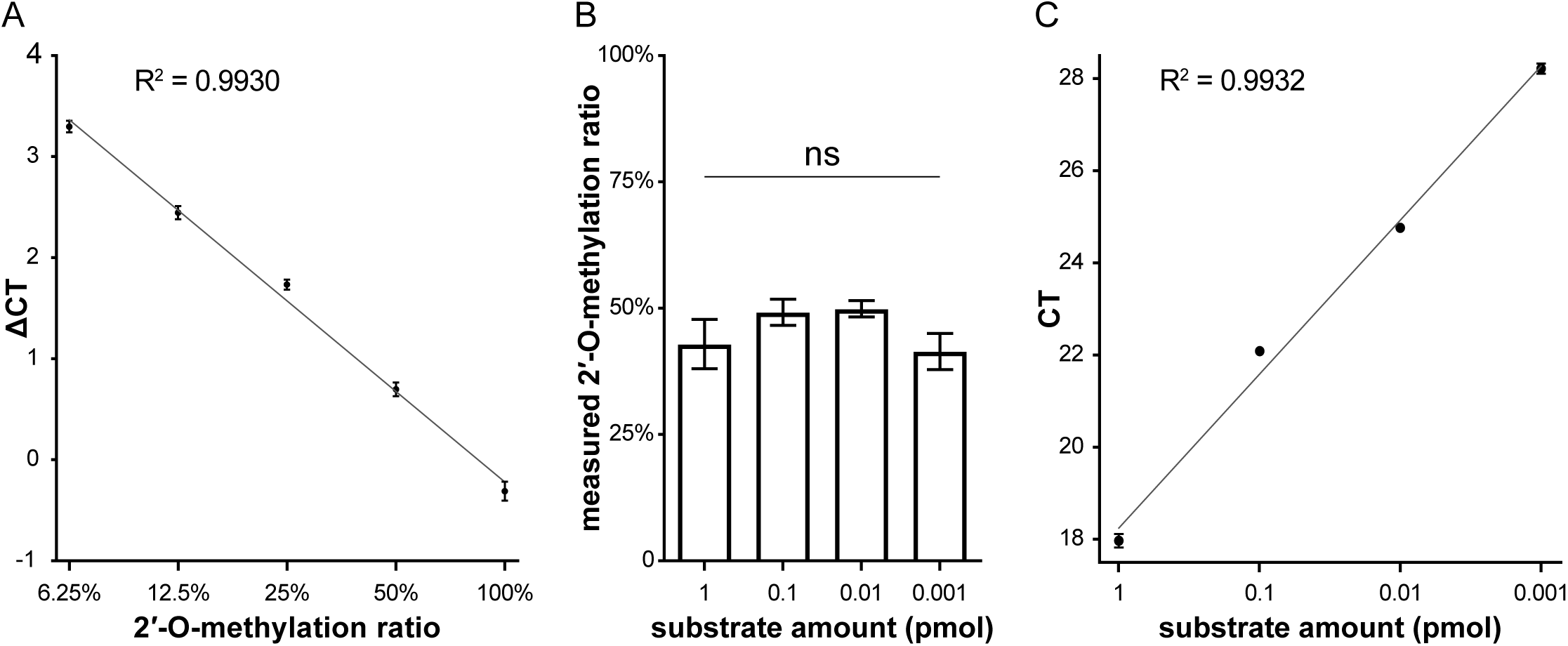
The construction of Nm-VAQ. (A) Correlation of Nm ratio of substrate to ΔCT for Nm-VAQ assay. RNase H-dependent site-specific cleavage and qRT-PCR were combined to form Nm-VAQ. The 0.1 pmol substrate with and without Nm were mixed to obtain a known Nm ratio, and ΔCT (Cycle Threshold) was from the RNase H treatment and the control samples. Error bars describe SD (n = 3). (B) Measure of the Nm ratio of four substrate amounts with known ratios of 50%. The Nm ratio was calculated by ΔCT values according to the above coefficient. Error bars describe SD (n = 3). ns, not significant, Brown-Forsythe ANOVA test. (C) Linear relationship between substrate amount and the product cleaved by RNase H. CT values were deprived from 3B.

### Quantitative detection of HeLa rRNA Nm status by Nm-VAQ

Now, we started to use Nm-VAQ to evaluate five sites in Hela rRNA, including four previously reported 2′-O-methylation sites, 18S 159Am, 354Um, 1391Cm (3), a newly discovered site 28S 4109Cm (11), and an unmethylated sites as a negative control, 18S 1197 G. Meanwhile, we collected HeLa cells from 4 different sources to observe whether these rRNA Nm sites were conserved. As demonstrated in the results, 18S 159A and 1391C were highly Nm-modified throughout different HeLa cell strains with 80-100% 2′-O-methylation ratio, consistent with other methods results(3,18). Interestingly, the 18S 354U methylation ratio was from 17.7% to 37.8%, which was detected as unmethylated by some previous reports while methylated by other tools(3). In addition, the newly found 28S 4109C was turned out to be 2′-O-methylated from 69.5%-84.1% ratio among different strains, which confirmed its 2′-O-methylation status (Figure 4). Finally, as a negative control, 18S 1197 G presented a barely detected signal, again proving the accuracy of Nm-VAQ (Figure 4).

**Figure 4.**
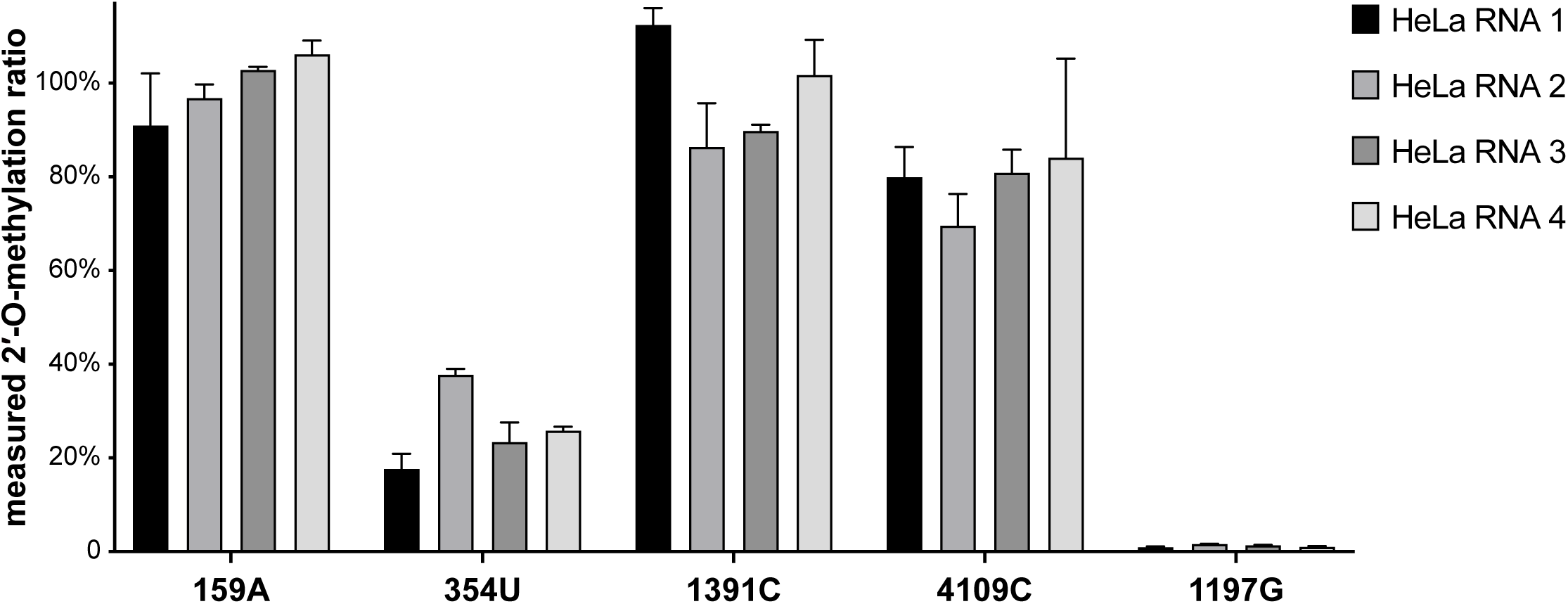
The Nm ratio of HeLa rRNA sites detecting by Nm-VAQ. 18s rRNA 159A, 354U, and 1391C sites were 2′-O-methylated reported by previous studies. 28s 4109C site was first discovered with Nm modification detected by NJU-seq. An unmethylated site at 18s 1197G was used as a negative control.

## Discussion

Several previously developed low-throughput Nm detection methods, including LC-MS, RTL-P, and DNA polymerase, have various defects. LC-MS is labor-intensive and difficult for mRNA Nm detection due to the requirement for many RNA molecules(13). Both RTL-P and DNA polymerase relied on the blocking of Nm on reverse transcription, which can be called RT-based methods(12,14). As the schematic illustrates, any Nm site between the amplification products will generate a methylation signal, and thus these two detections are non-site-specific methods (Supplementary Figure 3). Meanwhile, although both RTL-P and DNA polymerase methods could acquire linear results correlated with methylation ratio with synthetic RNA, the result varied with different amounts of target RNA. In the RTL-P method, RT-fold change is negatively correlated with the Nm amount, but cannot indicate the absolute proportion of the modification(14). In addition, the original study of RTL-P mentioned the false positive and negative results, which may be caused by RNA secondary structure that can occur on several rRNA sites(14). Compared to those methods, Nm-VAQ demonstrated apparent advantages. Nm-VAQ anchored the cleavage position to target site directing by chimera probe and discriminated 2′-O-methylated RNA from unmethylated RNA molecules by RNase H. This method acquired the absolute amount of the accurate 2′-O-methylation of target site simultaneously. In addition, Nm-VAQ showed its capability to consistently evaluate targets with low amounts or low methylation ratios, which was critical for the study of mRNA and other RNAs.

Our study systematically explores how RNase H, chimera probe, and substrate determined the cleavage site. By testing different chimera structures of DNA and RNA combination, we concluded to anchor the RNase H cleavage site with D_(4)_R_(13)_-(Nm). It was interesting to see RNase H prefer to cleave RNA substrate 4 nt upstream from the DNA-RNA boundary of the chimera probe. In future studies, the cleavage molecular mechanism of such unnatural hybrid nucleotides might be explained by a co-crystal structure of RNA substrate, chimera probe, and inactivated RNase H protein.

The results of HeLa cell were worth further exploration. With the help of Nm-VAQ, the 2′-O-methylation status of HeLa rRNA sites was not consistent. For example, 18S 1391C showed ∼100% 2′-O-methylation in HeLa cell 1, but only ∼80% in HeLa cell 2 and 3; 18S 354U showed ∼20% 2′-O-methylation in HeLa cell 1, but ∼40% in HeLa cell 2. As reported in several recent articles, the 2′-O-methylation status of HeLa rRNA sites varied due to strain difference, growth conditions, and genomic instability(18,19). Although the role of various modification status of these Nm sites is unknown, it may contribute to ribosome population heterogeneity further to impact translation. Nm-VAQ provided a solution to access the study of this direction.

---

**Supplementary Figure 1.**
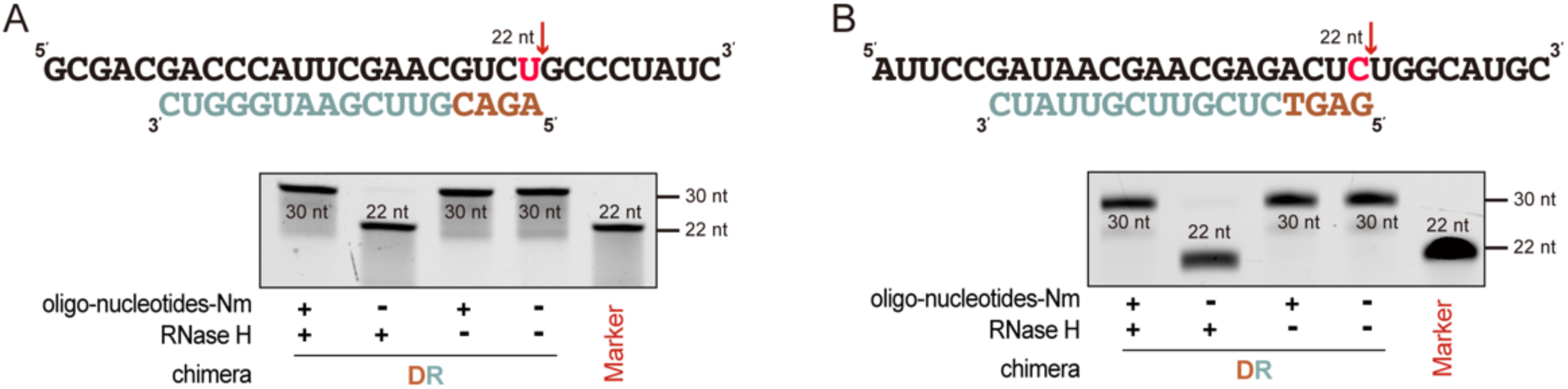
The RNase H cleavage guided by the DR chimera probe was inhibited by 2′-O-methylation. (A, B) The cleavage of RNase H in the other two oligo-nucleotides. The red sites indicated the Nm site, and the red arrows indicated that the cleavage sites. The number presented the length of FAM-labeled cleavage products.

**Supplementary Figure 2.**
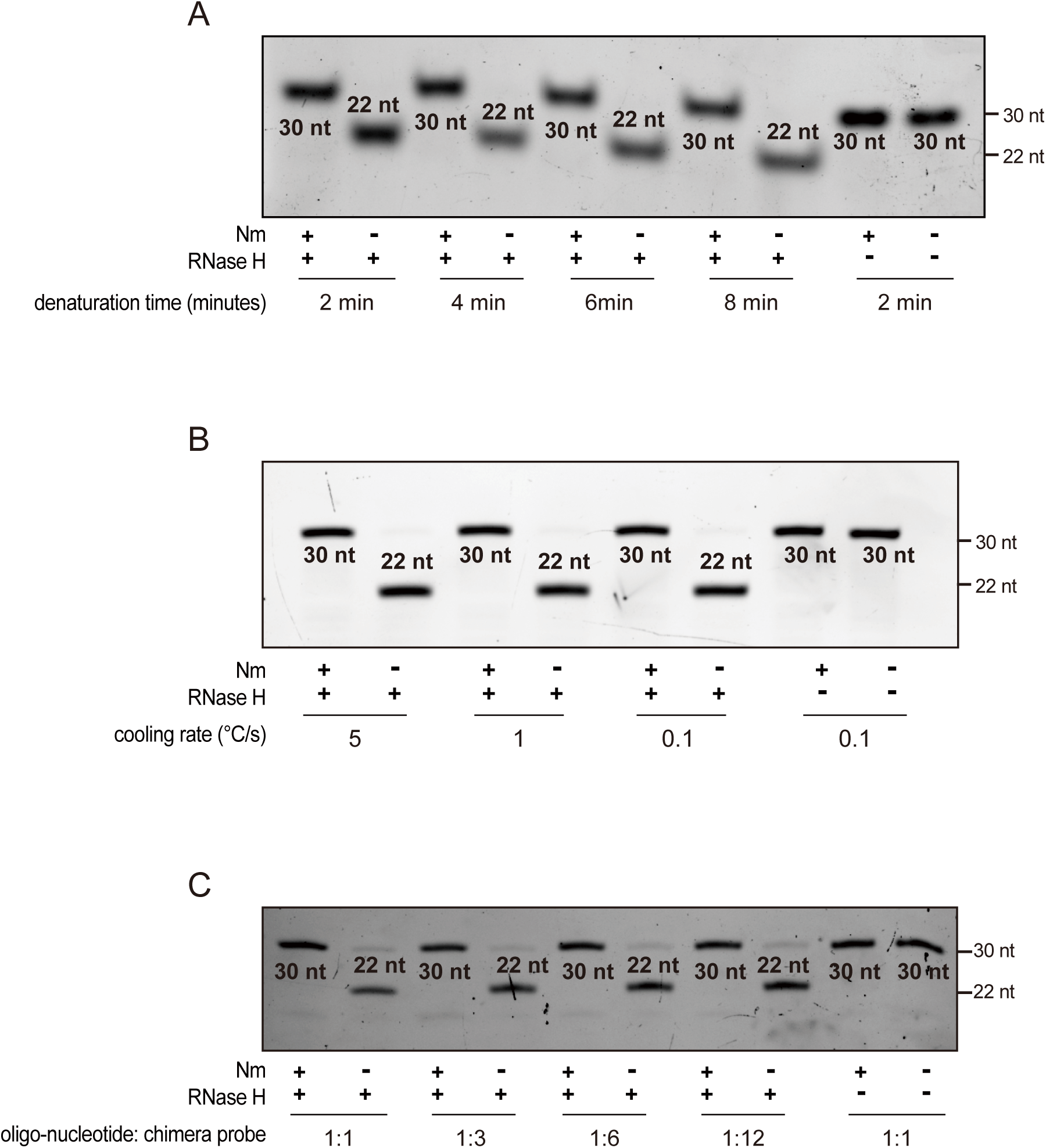
Exploration of RNase H hydrolysis condition. (A) RNase H cleavage with different denaturation time at 95°C. (B) RNase H cleavage with different cooling rate from 95 °C to 22 °C. (C) RNase H cleavage with a different molar ratio of oligo-nucleotide and chimera probe.

**Supplementary Figure 3.**
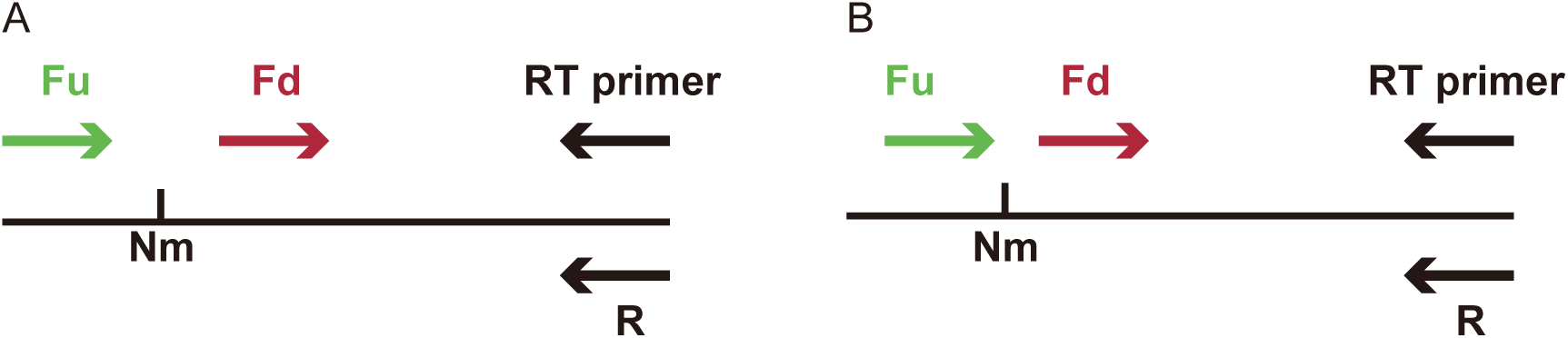
The schematic of the RT-based methods. (A) The 2′-O-methylation detection of the RTL-P method(14). The RT (reverse transcription) primer is positioned downstream of Nm site. Two forward primers for the subsequent PCR amplification were designed located either downstream (Fd) or upstream (Fu) of the Nm site. The RT primer was used to be the R primer in PCR amplification. (B) The 2′-O-methylation detection with an engineered DNA polymerase(12). The 3′ end of Fu primer was located on 1nt upstream of Nm site, and the 5′ end of Fd primer was 5-6nt downstream of Fu primer.

